# HUMAN BREAST TUMOUR CELLS VIABILITY EFFECT OF AFRICAN DIOSCOREA ROTUNDATA TUBER EXTRACTS IN MCF-7 AND MDA-MB231 CELL LINES

**DOI:** 10.1101/2020.05.08.084269

**Authors:** Joy Ifunanya Odimegwu, Olukemi Abiodun Odukoya, Alejandro Español, Maria Elena Sales

## Abstract

**Objective:** We aim to test the efficacy of edible *Dioscorea species* grown and consumed in Nigeria, Africa on two breast cancer cell lines; MCF-7 and MDA-MB231 derived from a luminal and a triple-negative breast cancer (TNBC) respectively and to confirm safety in non-tumour cells MCF-10A using a well established cytotoxic compound paclitaxel as a standard. Metastatic breast cancer is a prevalent cause of mortality in women around the world. Breast cancer therapies have greatly advanced in recent years, but many patients develop cancer re-occurrence and metastasis and subsequently yield to the disease because of chemoresistance.

**Methods:** Ethanolic extracts of *Dioscorea rotundata* boiled and raw (DiosB and DiosR) respectively were chemically analysed for the presence of diosgenin using HPLC and the cytotoxic activity of the extracts were tested on MCF-7, MDA-MB-231 and MCF-10A cells *In vitro* by MTT assay.

**Results:** DiosB and DiosR extracts showed a higher maximal effect on MCF-7 cells than on MDA-MB231 after 24 h and 48 h treatments (p<0.0001 and p<0.05 respectively). DiosR, if applied at a range between 50-70 g/ml, can be effective to reduce breast tumor cell viability without affecting non tumorigenic MCF-10A cells either at 24 h or at 48 h. DiosB showed an IC_50_ of 38.83μg/ml while DiosR showed an IC_50_ of 41.80μg/ml.

**Conclusion:** These results show that ethanolic extracts of *Dioscorea rotundata* tubers could be used effectively to treat breast cancer tumors and this is in sync with its diosgenin content as other *Dioscorea species* applied for similar treatments in Asia and elsewhere.

## 1.0 INTRODUCTION

*Dioscorea rotundata* Poir. is a synonym of *Dioscorea cayennensis* subsp. *rotundata* (Poir.) J.Miège (1and 2) also called white yam (3 and 4) Fig. 1, A, B, C *Dioscorea rotundata* of the family Dioscoreaceae is a high value but an under-utilized, under-researched crop as millions of people in developing countries depend on the tubers as a source of staple food because of its high starch content and on its sales for livelihood. There is between 70% to 80% of dried matter of white yams are starch (5). About 97% of total world production and consumption of yams in general occur in Africa (6). Nigeria, in West Africa is the greatest producer of yams yielding more than 64% of the global crop total. (7).

**Fig 1.**
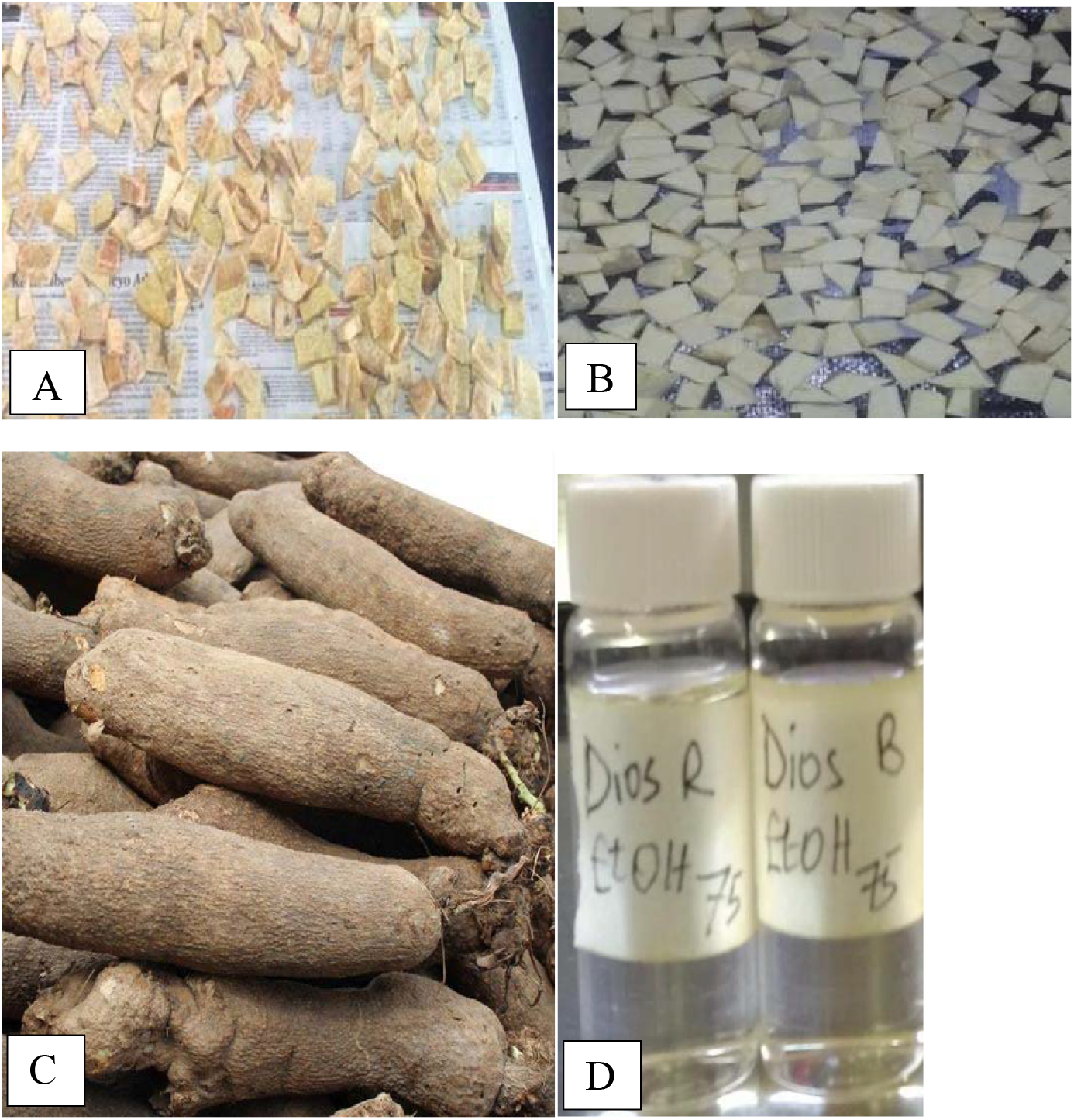
A. Boiled minisettes of white yam (DiosB). B. Raw minisettes of white yam (DiosR) C. White yam tubers on display at a market. D. Ethanolic extracts of DiosB and DiosR.

Diverse *Dioscorea* species have been in use in traditional Chinese medicine; *D. bulbifer*a has anti-tumour actions (8). Decoctions from yam leaves and tubers are used by traditional medicine to regulate women’s fertility, relieving painful periods and peri- and post-menopausal symptoms (9). It is also reported that wild yams might reduce the risk of breast cancer and cardiovascular diseases in post-menopausal women (10). *D. villosa* and *D. bulbifera* are the most widely used as drugs in Asia (11) most probably because of their diosgenin content. In Africa, *D. alata*; greater yam and *D. rotundata;* white yam are more common and preferred for consumption than others (Mignouna *et al.*, 2003), it is used medicinally traditionally as *D.cayenensis* its synonym due to its high antioxidant content against oxidative damage that could lead to cancer (4) and as *D. rotundata* with appreciable diosgenin content which has chemo-preventive action against chronic inflammation associated with cancer cells (3). Yams, like other plants can synthesize and accumulate a great deal of phytochemicals in their roots, leaves and vines as a part of its overall defence strategy against pests and diseases in vegetable kingdom. These phytochemicals are known to be useful in human illness. In previously reported research work on yam’s phytochemicals, it has been proved that tubers contain a glycosidic form of steroidal saponin/sapogenin which can be cleaved by acidification into the sugar moiety and diosgenin. Steroidal extracts from *D. villosa*, *D. floribunda* and *D. deltoidea* are used to derive sapogenin and diosgenin used by pharmaceutical companies in the manufacture of cortisone, and other steroids (12). Edible Dioscorea species; *D. rotundata* have not been studied for their phytochemical contents and medicinal values which could be immense and impactful (13, 14 and 15). Steroidal saponins are less widely distributed in the plant kingdom, compared to terpenoids, although their chemical structures have been useful for the development of pharmaceuticals with steroidal activities such as diosgenin from yam. Steroidal saponin-containing plants have therefore been of greater interest for use as phytopharmaceuticals, rather than as nutraceuticals (16).

Gynaecological malignancies are one of the leading causes of death worldwide in women. Epidemiologically, breast cancer is the most frequent type of tumor in women in both developed and developing countries. The incidence of breast cancer is increasing in the developing world due to increased life expectancy, increased urbanization and the adoption of Western lifestyles. Traditional chemotherapy has been partially useful to treat breast cancer patients because, although it can reduce tumor growth, concomitantly with undesirable effects and chemoresistance. Development of new treatment strategies that reduce costs and adverse effects are recommendable; for these reasons natural products, with demonstrated anti-tumor actions appear as a better treatment alternative. Like most epithelial tumors, breast cancer is a complex disease due to intra-tumor and inter-tumor heterogeneity. Breast tumors can be classified into 4 groups according to their genetic profile: Luminal A, express estrogen receptor (ER)+, Luminal B are ER+ and/or progesterone receptor (PR)+, and HER2+ or HER2-; and Basal (or triple negative) (ER− / PR− / HER2−). The survival of patients, after 5 years, decreases significantly from [1] to [4]. The decreasing percentage of survival is related to increased resistance to chemotherapy and/or radiotherapy and to a greater likelihood of relapse and/or metastasis formation. MCF-7 and MDA-MB231 cells represent the two extremes of this classification. MCF-7 cells derived from a human breast luminal A adenocarcinoma while MDA-MB231 cell line was obtained from a highly aggressive, invasive and poorly differentiated triple-negative breast cancer (TNBC) (17).

The purpose of this research is to carry out phytochemical assays on edible *D. rotundata* and to analyze the anti-tumour actions of two different extracts on human breast tumor cells and to compare them with the conventional chemotherapeutic agent paclitaxel.

## 2.0 METHODOLOGY

### 2.1 Plant materials

The tubers of *Dioscorea* species were obtained from International Institute of Tropical Agriculture, IITA Ibadan, *D. rotundata* (TDR 3731) and from a vendor at Ojuwoye market of Mushin Local Government Area, Lagos State. Nigeria.

### 2.2 Preparation of *D. rotundata* samples

The tubers were peeled, cut up into minisettes and washed. A portion of minisettes were boiled (DiosB) in clean water in an aluminium pot for 15 minutes and then strained and dried while the other half was also peeled, washed and cut into raw minisettes (DiosR). They were air dried during 72 h and then oven dried at 60°C for 15 min then milled into coarsely powdered samples and stored in glass bottles until needed.

### 2.3 Extraction and Thin-layer chromatography of DiosB and DiosR extracts

DiosB and DiosR samples each were weighed and extracted by cold maceration. The powdered samples were extracted in aqueous ethanol (25:75) by cold maceration three times and then filtered and dried in open air. The recovered dry extracts were saved at 4°C until used. Preparative TLC was carried out on silica gel plates (Kieselgel G, F254, type 60, Merck, Germany) eluted with Chloroform, Toluene, Methanol, Ethyl acetate in 7: 2.4: 0.8: 0.4, Diosgenin was used as standard. The developed TLC plate was observed under UV at 254 nm and 366nm. Visualization was also further achieved by spraying with Distilled water in chilled acetone, 60% Perchloric acid and Anesaldehyde in the following proportions 8:2:1:0.5 heat was applied in an oven at about 110 °C for 3–5 min. Plates were scanned or photographed while color of spots were still visible.

### 2.4 Extraction of (TDR 3731) tubers using Soxhlet apparatus

Dried and powdered (TDR 3731) tubers were packed into the thimble and then inserted into the Soxhlet extractor. The Soxhlet was inserted into the quick fit round bottom flask containing n-hexane with temperature of 40-60O C. The obtained extracts were subjected to Thin Layer Chromatography (TLC) and Gas Chromatography-Mass Spectrometry (GC-MS).

### 2.5 GC-MS analysis of (TDR 3731) tuber hexane extracts

A sample of TDR 3731 was injected into the Agilent technologies 7890GC-MS machine with number G3170-80026. The oven temperature program ranged from 70°C to 250°C, programmed at 3°C/min, with initial and final hold time of 2 min, carrier gas: He at 10 psi constant pressure, a split ratio of 1 : 40; injector, transfer line, and source temperatures were kept at 250°C; ionization energy 70 eV; mass scan range 40–450 amu. Characterization was achieved on the basis of retention time, elution order, in GC-FID capillary column (Aldrich and Fluka), mass spectra library search (NIST/EPA/NIH version 2.1, and Wiley registry of mass spectral data 7^th^ edition).

### 2.6 High Performance Liquid Chromatography Analysis of DiosB and DiosR

For Calibration Curve and Linearity studies; The specific system used is a combination of Waters HPLC System and Agilent 1100 HPLC with a quartenary pump, an auto liquid sampler (ALS), a thermostated column compartment and a variable wavelength detector (VWD). The analysis is carried out at a constant flow rate of 1.0 ml/ min all throughout. The mobile phase is a mixture of acetonitrile and water in a ratio 90:10 v/v. for diosgenin. The different components in our samples are detected by a variable wavelength UV detector set at 203 nm and the data station displays the recorded data in the form of chromatograms at volumes 20μL. The extracts were duly prepared for HPLC. Calibration curve was obtained from the standards working solutions for linearity studies and was plotted using Microsoft Excel. Every sample, from the standards to the *Dioscorea* samples, were prepared and injected into the HPLC system.

### 2.7 Cell culture

The human breast adenocarcinoma cell line MDA-MB231 (CRM-HTB-26) and MCF-7 (HTB-22) were obtained from the American Type Culture Collection (ATCC; Manassas, USA) and cultured in DMEM (Invitrogen Inc., Carlsbad, USA) with 2 mM L-glutamine and 80 μg/ml gentamycin, supplemented with 10 % heat inactivated fetal bovine serum (FBS) (Internegocios SA, Mercedes, Argentine) at 37°C in a humidified 5% CO_2_ air. MCF-10A cells (CRL-10317) were also purchased by ATCC and constitute a non-tumorigenic cell line derived from human mammary tissue. These cells were grown on tissue culture plastic dishes in DMEM:F12(1:1) (Invitrogen Inc., Carlsbad, USA) supplemented with 10% FBS, hydrocortisone (0.5 μg/ml), insulin (10 μg/ml), and hEGF (20 ng/ml). Cell lines were detached using the following buffer: 0.25% trypsin and 0.02% EDTA in Ca2+ and Mg2+ free PBS from confluent monolayers. The medium was replaced three times a week. Cell viability was assayed by Trypan blue exclusion test and the absence of mycoplasma was confirmed by Hoechst staining.

### 2.8 MTT assay

Cell viability was evaluated by a colorimetric assay using the reagent 3- (4,5-dimethylthiazol-2-yl)-2,5-diphenyl tetrazolium bromide (MTT) Cells were seeded in 96-well plates (10^4^ cells/well) in 200 μl of DMEM culture medium supplemented with 5% of FBS at 37° C in a humidified atmosphere with 5% CO_2_. The cells were deprived of FBS for 24 h and then treated with increasing concentrations of DiosB, DiosR, diosgenin and paclitaxel (the latter 2 were used as controls to induce cell death) added during 24 h or 48 h. After treatment, medium was replaced by fresh DMEM medium without phenol red, and MTT solution (0.5% in PBS) was added. Finally the absorbance at 540 nm was measured in an ELISA reader computerized with the Gen5 program and the cell viability was calculated as a percentage with respect to the control (untreated cells considered as 100%).

## 3.0 RESULTS AND DISCUSSION

Thin layer Chromatography showed presence of diosgenin in hexane extract of TDR (3731) and ethanolic extracts of DiosB and DiosR (Fig. 1 D) confirmed by HPLC. This is very interesting considering that previous works on Dioscorea analyzing the content of diosgenin had been performed on inedible species like *D. zingiberensis, D. septemloba, D. collettii* (18). Kanu *et al*., 2018 mentioned the presence of diosgenin in white yam in an inconclusive manner as they were unable to confirm diosgenin presence with chemical analysis. The discovery of diosgenin in edible yams is huge as the tubers are staple foods to hundreds of thousands of indigenious people in Africa and particularly in Nigeria. Also Caryophylene and Pthallic acid were discovered in the hexane extracts of TDR (3731) from the GC-MS analysis (Appendix 1).

Diosgenin is present in both DiosB and DiosR as it was confirmed by HPLC, Fig. 3 and 4. We observed that the amount of diosgenin in the boiled sample is higher than that in the raw one Figures 3 and 4 Edible *Dioscorea* cultivated in Africa has been never analyzed regarding diosgenin content and for this reason our findings could constitute novel and interesting knowledge for botanical medicine. Chinese species of Dioscorea; *D. zingiberesis, D. oppositifolia and D. japonica* have diosgenin (18, 23, 25).

**Figure. 2.**
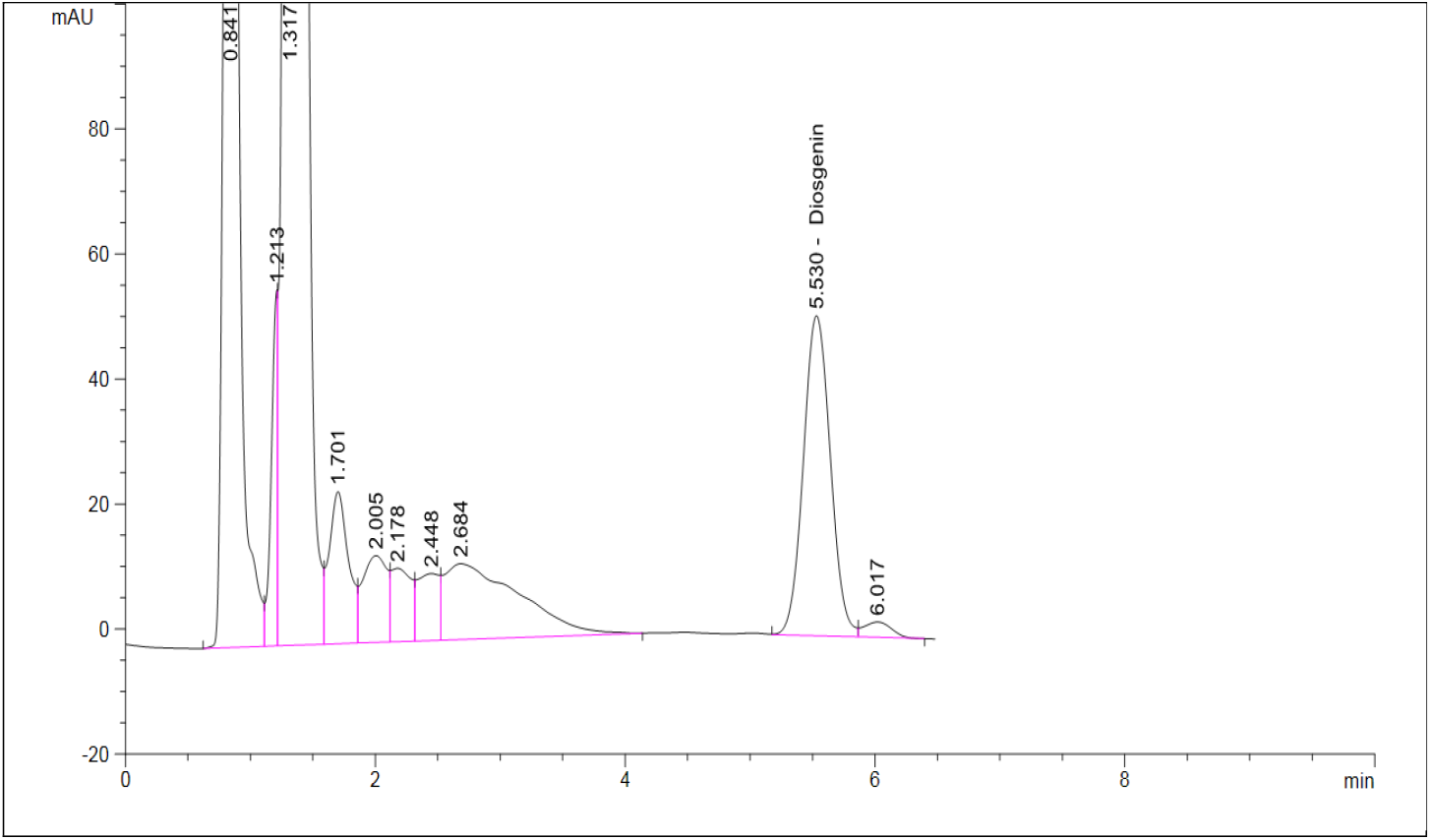
**HPLC** Chromatogram of diosgenin.

**Figure 3:**
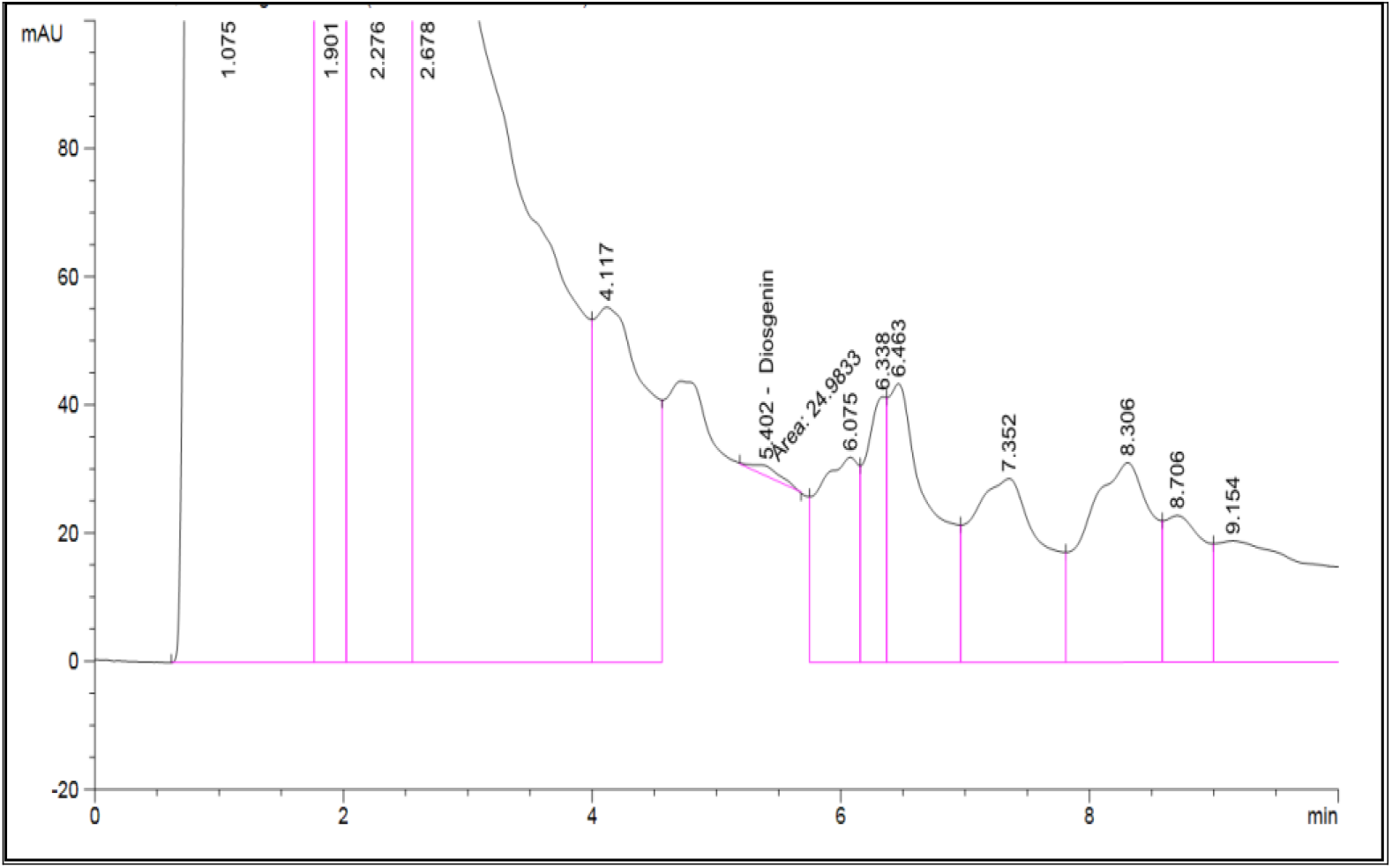
**HPLC** Chromatogram of DiosR showing diosgenin peak

**Figure 4:**
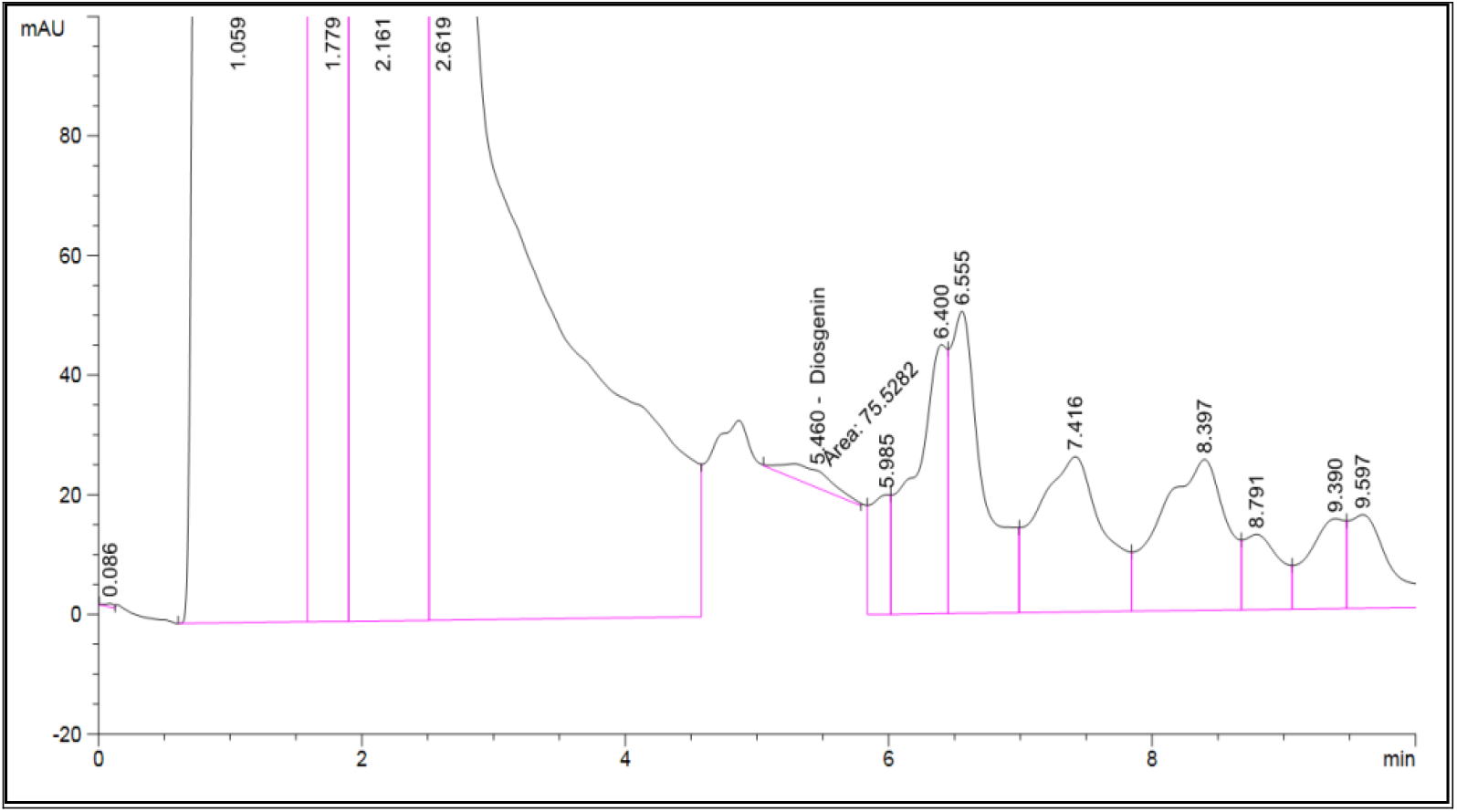
Diosgenin chromatogram of DiosB showing diosgenin peak

Diosgenin has already been reported to exert anti-proliferative effects on different types of cancer cells including breast tumor cells (19, 24, 26). Here, we confirmed these actions on MDA-MB231 and MCF-7 cells obtaining concentration-dependent actions after 24 h and 48 h treatment. Unfortunately, diosgenin was also effective and similarly potent on non-tumorigenic MCF-10A cells as an undesirable action, Here, we analyzed the action of DiosR and DiosB that contain diosgenin on tumor cells. We observed that, both extracts were effective to reduce tumor cell viability in a concentration-dependent manner but in a less potent manner than pure diosgenin. No differences were observed between both extracts and both times of treatment on MCF7 cells (Fig. 5A and B); but either DiosB or DiosR exerted a time dependent action on MDA-MB231 cells being more effective at 48 h than at 24 h (Fig 6). In spite of this, both extracts were more effective on MCF-7 than on MDA-MB231 cells showing a higher maximal effect on MCF-7 cells than on MDA-MB231 either after 24 h or 48 h of treatment (p<0.0001 and p<0.05 respectively) probably due to the high aggressiveness and invasiveness documented by many authors for triple negative MDA-MB231 tumor cells (20, 21).

**Figure 5:**
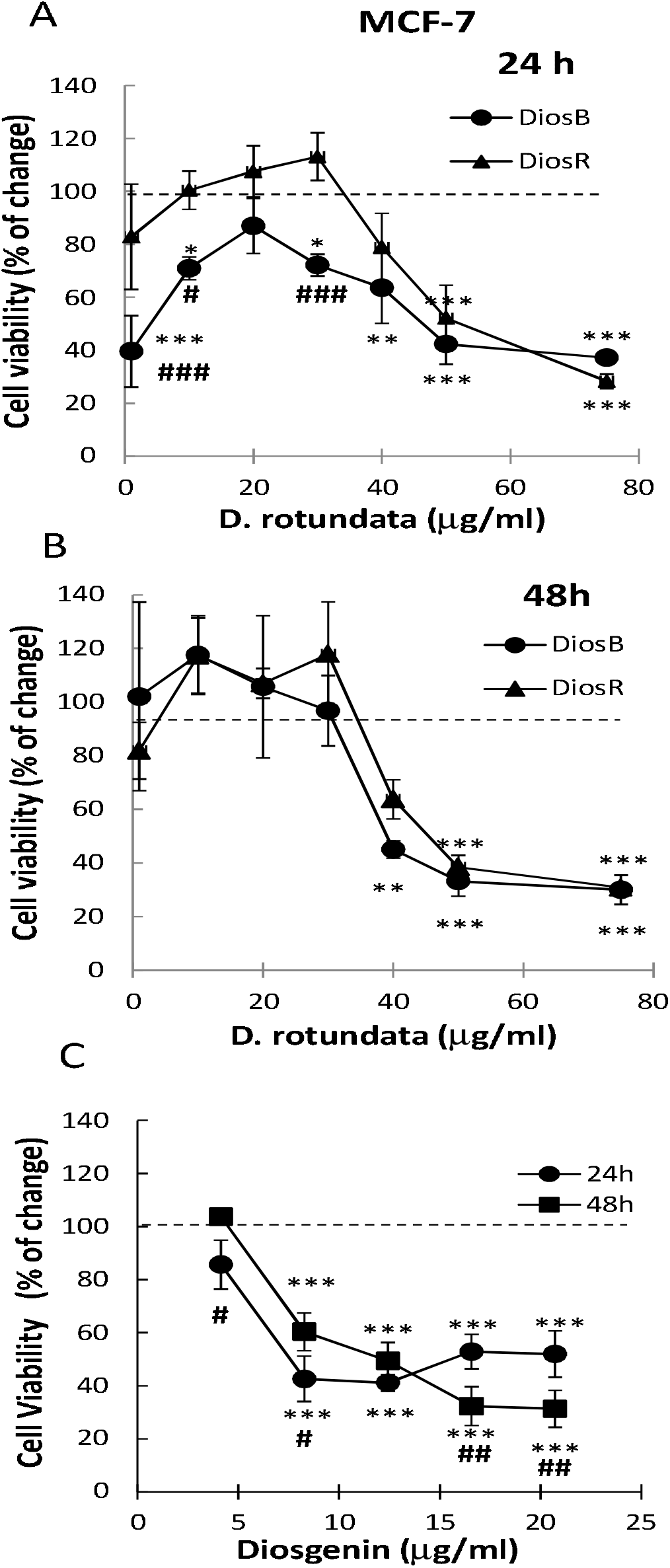
Modulation of MCF-7 breast tumor cell viability. Concentration-response curves of *D. rotundata* extracts boiled (DiosB) or raw (DiosR) added at A) 24 h, B) 48 h and C) concentration-response curves of Diosgenin added during at 24 h and 48 h. Results were expressed as percentage of change respect to control (cells without treatment considered as 100%). Values are mean±S.E.M. of 4 experiments performed in quadruplicate (#P<0.05; ##P<0.01; ###P<0.0001 DiosB vs. DiosR. *P<0.05; **P<0.001; ***P<0.0001 vs. control.

**Figure 6:**
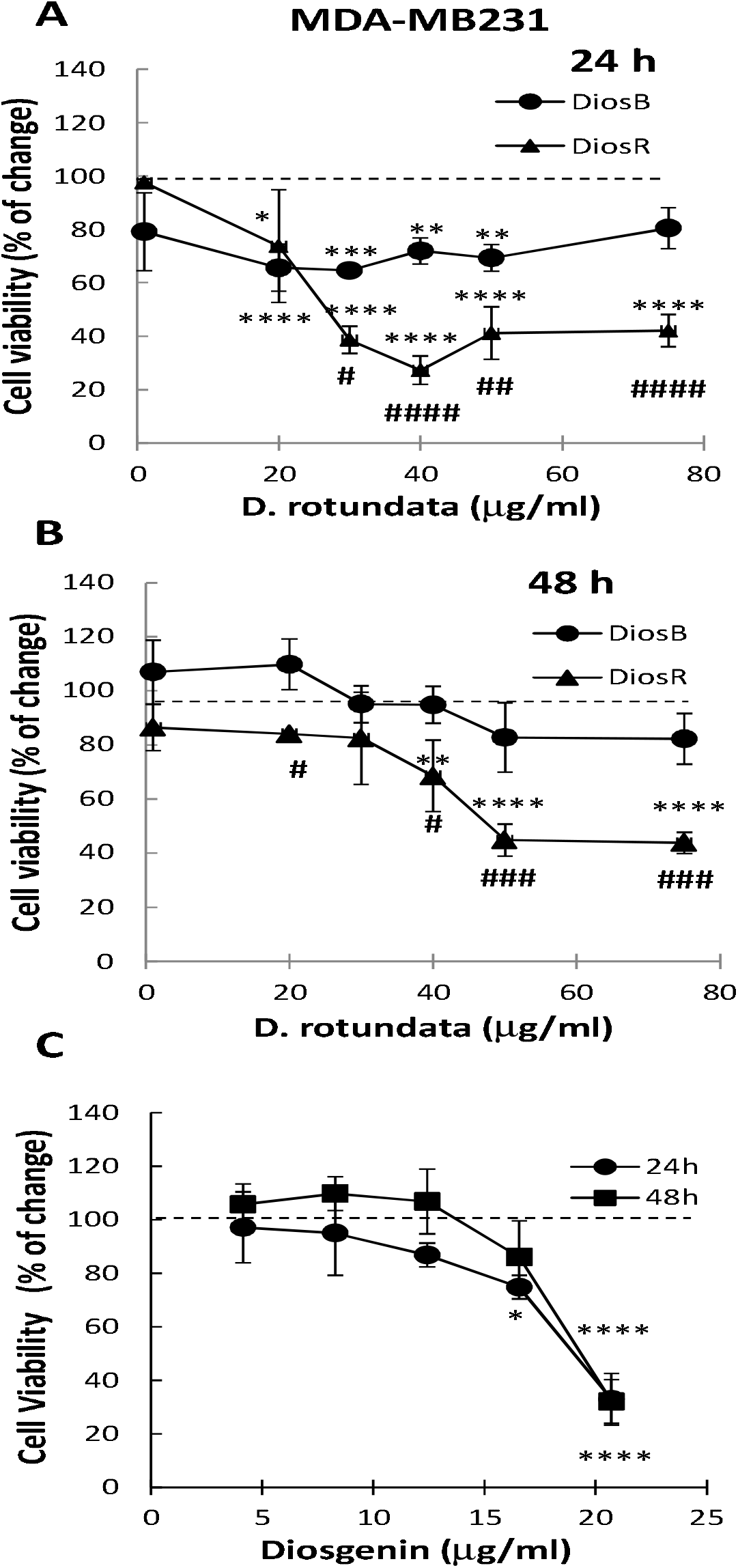
Modulation of MDA-MB231 breast tumor cell viability. Concentration-response curves of *D. rotundata* extracts boiled (DiosB) or raw (DiosR) added at A) 24 h, B) 48 h and C) concentration-response curves of Diosgenin. Results were expressed as percentage of change respect to control (cells without treatment, considered as 100%). Values are mean±S.E.M. of 4 experiments performed in cuadruplicate. (#P<0.05; ##P<0.01; ###P<0.001; ####P<0.0001 DiosB vs. DiosR. *P<0.05; **P<0.01; ***P<0.001; ****P<0.0001 vs. control).

Another important finding of our work is that both extracts, but mainly DiosR were more effective on tumor cells than in non-tumor MCF-10A human breast cells (Fig. 7 and Table I). This could be taken as an advantage in comparison to traditional chemotherapy. It is important to note that if DiosR is used at a range between 50-70 g/ml it can be effective to reduce breast tumor cell viability without affecting non tumorigenic MCF-10A cells either at 24 h or at 48 h (Fig. 5-7).

**Figure 7:**
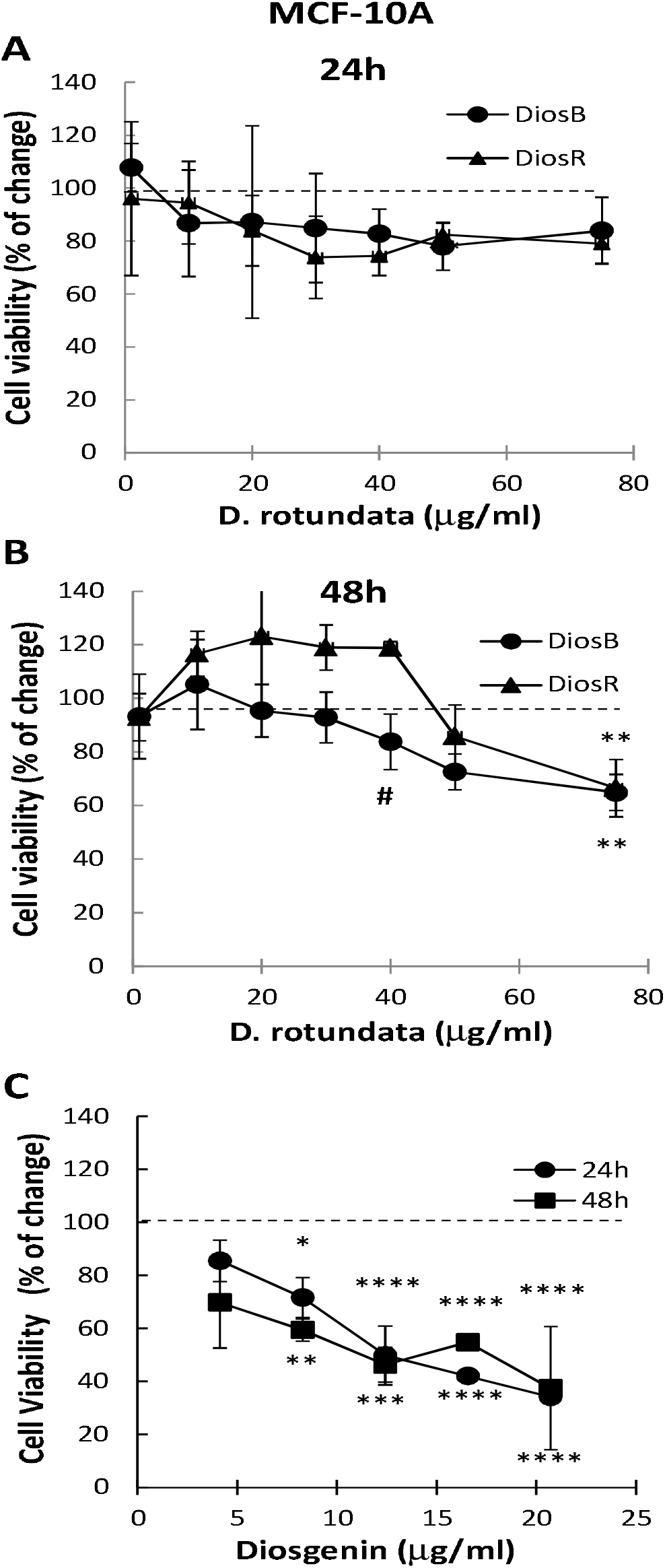
Modulation of MCF-10A non-tumorigenic breast cell viability. Concentration-response curves of *D. rotundata* extracts boiled (DiosB) and raw (DiosR) added during A) 24 h, B) 48h and C) concentration-response curves of Diosgenin added during 24 and 48h. Results were expressed as percentage of change respect to control (cells without treatment considered as 100%). Values are mean±S.E.M. of 4 experiments performed in quadruplicate. (#P<0.01 DiosB vs. DiosR. *P<0.05; **P<0.01; ***P<0.001; ****P<0.0001 vs. control).

**Table I.** Comparison of the effect exerted by DiosR on human breast cells.

| **Cell Line** | **DiosR**<br>(75 μg/ml) | |
| --- | --- | --- |
|  | **Time**<br>(h) | **Effect**<br>(% of inhibition) |
| **MCF-7** | 24 | 72±3<sup>ac</sup> |
| | 48 | 69±1<sup>bd</sup> |
| **MDA-MB231** | 24 | 58±6<sup>ac</sup> |
| | 48 | 56±4<sup>bd</sup> |
| **MCF-10A** | 24 | 21±7<sup>a</sup> |
| | 48 | 34±10<sup>b</sup> |

Cells were treated with the maximal effective concentration of DiosR (75 μg/ml) during 24 and 48 h. Results were expressed as percentage of inhibition in cell viability with respect to control (cells without treatment considered as 100%). Values are mean ± S.E.M. of 4 experiments performed in quadruplicate. MCF-7 and MDA-MB231 cells vs. MCF-10A cells 24 h (a) $p<0.0001$ ; 48 h (b) $p<0.0001$ ; MCF-7 cells vs. MDA-MB231 cells 24 h (c) $p<0.05$ and 48 h (d) $p<0.05$ .

As shown in Table II, lower concentrations of DiosR are needed to reduce the viability of tumor cells in comparison to normal cells by 25%, similarly to paclitaxel, a chemotherapeutic agent obtained from the plant, Pacific yew, *Taxus brevifolia* frequently used to treat breast cancer patients (22), revealing a certain degree of specificity for DiosR as an anti-tumor compound.

**Table II.** Comparison of the effect of DiosR with paclitaxel.

| Cell Line | EC25 (48 h) |  |
| --- | --- | --- |
|  | DiosR ( g/ml) | Paclitaxel (M) |
| **MCF-7** | 38.79 | $3.97 \times 10^{-8} \text{M}$ |
| **MDA-MB231** | 37.97 | $5.95 \times 10^{-8} \text{M}$ |
| **MCF-10A** | 48.73 | $4.42 \times 10^{-7} \text{M}$ |

Cells were treated at 48 h with increasing concentrations of both compounds and the effective concentrations necessary to reduce to 25% cell viability (EC25) was calculated. Values are mean ± S.E.M. of 4 experiments performed in quadruplicate.

In conclusion our work demonstrates the effectiveness of two extracts obtained from *Dioscorea rotundata* that are able to reduce breast tumor cells viability derived not only from a Luminal A tumor, like MCF-7 cells but also from a triple negative breast tumor, MDA-MB 231 that is considered one of the most aggressive in the molecular classification of tumors. The administration of *Dioscorea rotundata* instead of conventional chemotherapeutic agents like paclitaxel could be a useful and cheaper strategy and also less harmful to treat breast cancer in developing countries employing natural medicine

## Acknowledgement

We wish to thank TETFUND, TWAS-UNESCO for the research fellowship offered to Dr. JI Odimegwu, and CEFYBO-CONICET for accommodation and laboratory financial support offered to Odimegwu, JI. Also we wish to thank members of the Laboratory of Tumor Immunopharmacology; Center for Pharmacological and Botanical Studies, Francisco and Martin among others for all their help.

